# High prevalence of SARS-CoV-2 antibodies in pets from COVID-19+ households

**DOI:** 10.1101/2020.09.22.307751

**Authors:** Matthieu Fritz, Béatrice Rosolen, Emilie Krafft, Pierre Becquart, Eric Elguero, Oxana Vratskikh, Solène Denolly, Bertrand Boson, Jessica Vanhomwegen, Meriadeg Ar Gouilh, Angeli Kodjo, Catherine Chirouze, Serge Rosolen, Vincent Legros, Eric M. Leroy

**Affiliations:** Institut de Recherches et de Développement (IRD), Maladies Infectieuses et vecteurs: Ecologie, génétique, Evolution et Contrôle (MIVEGEC) (IRD 224 - CNRS 5290 - UM), Montpellier, France (; Service de maladies infectieuses et tropicales, CHRU hôpital Jean-Minjoz, Besançon, France; Université de Lyon, VetAgro Sup, Campus vétérinaire de Lyon, Marcy-l’Etoile, France; Environment and Infectious Risk Unit, Laboratory for Urgent Response to Biological Threats, Institut Pasteur, Paris, France; CIRI – Centre International de Recherche en Infectiologie, Team EVIR, Univ Lyon, Université Claude Bernard Lyon 1, Inserm, U111, CNRS, UMR5308, ENS Lyon, Lyon, France; The OIE Collaborating Centre for Detection and Identification in Humans of Emerging Animal Pathogens, Institut Pasteur, Paris, France; Groupe de Recherche sur l’Adaptation Microbienne (GRAM 2.0), Normandie Univ, UNICAEN, UNIROUEN, EA2656, Caen, France; Laboratoire de Virologie, Centre Hospitalo-Universitaire, Caen, France; UMR Chrono-Environnement - Université de Bourgogne Franche-Comté, Besançon, France; Sorbonne Université, INSERM, CNRS, Institut de la Vision, Paris, France; Clinique Veterinaire Voltaire, Asnieres, France

**Author notes:** Drs. Fritz and Rosolen contributed equally to this article. Drs. Krafft and Becquart contributed equally to this article. Drs. Legros and Leroy contributed equally to this article. **Corresponding author: Vincent Legros:** CIRI, 46 allée d’Italie 69007 Lyon, France; E-mail address, **Eric M.Leroy :** UMR IRD-CNRS-UM, 911 avenue Agropolis, 34394 Montpellier, France.

## Abstract

In a survey of household cats and dogs of laboratory-confirmed COVID-19 patients, we found a high seroprevalence of SARS-CoV-2 antibodies, ranging from 21% to 53%, depending on the positivity criteria chosen. Seropositivity was significantly greater among pets from COVID-19+ households compared to those with owners of unknown status. Our results highlight the potential role of pets in the spread of the epidemic.

---

Since its emergence in December 2019, in Wuhan, China, Respiratory Syndrome Coronavirus 2 (SARS-CoV-2) has spread throughout the world, probably exclusively through human-to-human transmission. However, the existence of hundreds of millions of companion animals living closely with humans raises the question of their susceptibility to infection and potential role in the outbreak. Cats and dogs are known to be infected by Alphacoronaviruses and Betacoronavirus (Feline CoVs, Canine CoVs)^1^, and thus may be susceptible to SARS-CoV-2, which also belongs to the Betacoronavirus group. In Europe, the prevalence of canine coronavirus infection is low^2^. Feline coronavirus prevalence is higher^3–5^, with typical seroprevalence ranging from 50% in healthy Swiss cats to 37% in Japan. Additionally, epidemiological, biological, and virological characteristics of coronaviruses, mainly based on Spike-protein plasticity, suggest species barriers to infection may be easily crossed. Thus, pet contamination by sick owners is not only likely but perhaps expected, given the numerous opportunities for spillover^6–8^. The observation of several cases of mild infections in dogs and cats of infected owners, and serological surveys of pet populations reporting infection rates ranging from 0% to 15,8% ^9–12^, highlight this risk. Yet, despite these observations, studies continue to suggest that the risk of contamination of pets by their owners is low and that the role of pets in the spread of the outbreak is trivial or nonexistent.

Presently there is no published study accurately assessing the contamination levels in household pets. Here we present results from a serological survey of pets conducted between May and June 2020 in two neighbouring regions of eastern France: Franche-Comté and Rhone-Alpes. Both regions had similar epidemiological characteristics and health management policies, with the first hospitalised deaths registered in March 2020 (https://www.gouvernement.fr/info-coronavirus/carte-et-donnees). The first group of pets, from the Franche-Comté region, were living in homes where at least one person expressed respiratory symptoms and tested positive for SARS-CoV-2 at the University Hospital of Besançon (COVID-19+ household group). The second group, from the Rhone-Alpes, were pets from households where exposure was unknown (unknown status household group). Lastly, we included a control group of animals sampled in 2018 and early 2019 before the outbreak, including hyperimmune sera from ten cats with feline infectious peritonitis virus (FIPV), (Control group). Inclusion FIPV-infected cat sera in the control group allows us to exclude possible cross-reactivity of antibodies generated in response to non-SARS-CoV-2 coronaviruses.

We combined four different tests based on two different techniques to ensure the greatest degree of specific-antibody detection. Three microsphere immunoassays (MIA) detected anti-SARS-CoV-2 IgGs produced in response to viral N, S1, or S2 proteins, and a retrovirus-based pseudoparticle assay detected SARS-CoV-2 neutralizing antibodies (Methods). Taking into account these two types of assays, animals were declared COVID-19 positive following a positive seroneutralization assay or if they were positive for all three MIA tests. This positivity criterion ensures 100% specificity, as none of the animals in the control group tested positive for the three MIAs or for seroneutralisation (Fig. 1a-d).

**Figure 1.**
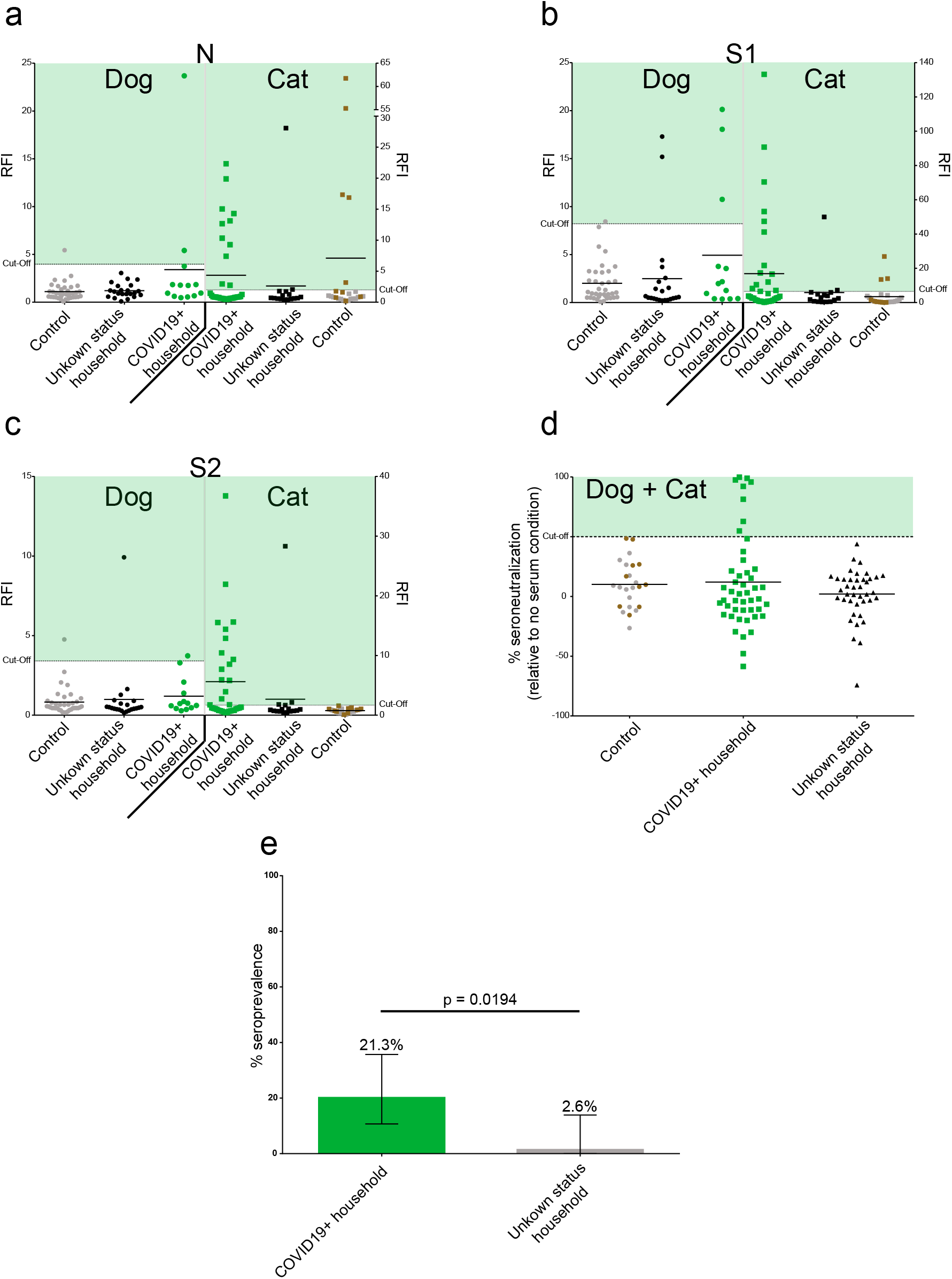
High prevalence of SARS-CoV-2 antibodies in COVID19+ household pets. Serological evaluation of anti-SARS-CoV-2 antibodies in pets from unknown status and COVID19+ households. COVID19+ households had at least one COVID-19 laboratory-confirmed person (Green). Unknown status households were those with no confirmed SARS-CoV-2 infected person (Black). Control include pre-pandemic population (Grey) and FIPV infected cats (Brown). **a**: Anti-N antibody levels. **b**: Anti-S1 antibody levels. **c**: Anti-S2 antibody levels. SARS-CoV-2 specific antibody levels were assessed using MIAs and expressed as Relative Fluorescence Intensities (RFI) to control antigen. A pre-pandemic population was used to determine the cut-off (mean + 3*standard deviation). **d**: Percentage of neutralizing activity in pet sera. Neutralising activity was assessed using a pseudoparticle assay and expressed as the percent neutralization relative to a no serum condition. For **a,b,c,d** mean line are presented. **e**: Prevalence based on positive anti-N, anti-S1, anti-S2, and seroneutralization tests in COVID19+ and unknown status households. 95% confidence interval are presented (± 95% confidence intervals).

A remarkably high 21.3% (10 of 47 animals tested) of pets in COVID-19+ households tested positive, including 23.5% of cats (8/34) and 15.4% of dogs (2/13), a non-significant difference (p=0.70) (Fig. 1a-e, Supplementary tables 1-2). Of the 16 cats and 22 dogs tested from households of unknown status, only one animal (a cat) tested positive (Fig. 1a-e, Supplementary tables 1-2), representing a significantly lower seroprevalence than the COVID-19+ group (p=0.0194). The risk of testing seropositive was eight times higher for pets sharing a home with a COVID-19+ person than for pets in homes of unknown status (relative risk of being seropositive = 8.1).

Though we cannot definitively prove that all the ten positive animals were infected with SARS-CoV-2, the much greater seroprevalence in animals from COVID-19+ households provides strong evidence that pets have been infected with SARS-CoV-2.

The highly variable antibody responses to SARS-CoV-2 in human infections^13,14^, calls into question our strict criteria for defining seropositive tests. If we consider an animal seropositive if any one test was positive, 53.2% in pets from COVID-19+ households show signs of having been infected (58.8% of cats (20/34) and 38.5% of dogs (5/13)) compared to 15.8% (6/38) of pets in homes of unknown status.

A recent Swiss study found that anti-N antibody assays substantially underestimate (i.e., by 30% to 45%) the proportion of SARS-CoV-2 exposed individuals compared to anti-S antibody assays in population-based seroprevalence studies^15^. Assuming similar dynamics in pets, the actual seropositivity in COVID-19+ households is likely closer to 53% than 21%, indicating that infection risk in the pets of COVID-19 positive owners is much higher than previously described. Given that cats and dogs may become infected, do they contribute to COVID-19 spread due to spillover back into humans? While viral shedding from pets does not appear sufficient for transmission to humans or other animals encountered during walks, for people in closer contact, precautionary measures should be considered as part of a ‘one health’ global control strategy.

## Methods

The dataset generated during the current study are available from the corresponding authors on reasonable request.

### COVID19+ household group

The COVID19+ household group was recruited from a cohort of 825 patients diagnosed with SARS-CoV-2 infection by reverse-transcriptase–polymerase-chain-reaction testing of nasopharyngeal swabs in the infectious tropical disease department at the University Hospital of Besançon between March 1 to April 25. From May 11 to 22, 384 patients were contacted and 84 reported owning dogs and/or cats. 34 gave us their informed consent to sample their pets. Whole blood samples were collected from 13 dogs and 34 cats between June 7 and June 12, 2 to 3 months after the owners were diagnosed.

### Unknown status household group

The unknown status household group recruited volunteers among staff and students at VetAgro Sup (Lyon’s National Veterinary School). Dogs and cats from all volunteers were included. The COVID-19 status of the pet owners was unknown. Blood samples were obtained from each animal (no selection) from 14th of May to 4th of June 2020. Clinical examination at the time of sampling indicated that all animals were healthy. Sampling of animals for this study was approved by VetAgro Sup ethical committee (approval number n°2031).

### Neutralization activity measurement

To measure the neutralizing activity in sera, we developed a MLV-based pseudoparticle carrying a GFP reporter pseudotyped with SARS-CoV2 spike (SARS-CoV-2pp). Briefly, SARS-CoV-2pp were incubated in 1/100 dilution of sera at 37°C for 1 hour. The mix was added on reporter cells (VeroE6), spinoculated for 2 hours (2.500g, 25°C). After 2 hours of incubation, the inoculum was removed and replaced with fresh medium and cells were incubated for 72h before FACS analysis. The level of infectivity was expressed as % of GFP positive cells and compared to cells infected with SARS-CoV-2pp incubated without serum. Prepandemic (including non SARS-CoV2 coronaviruses positive) sera from France were used as negative controls, and anti-SARS-CoV-2 RBD antibody was used as positive control. Seroneutralization specificity was 100% as already described.

### Microsphere immunoassay

Dog and cat serum samples were tested using a multiplex Microsphere immunoassay (MIA). 10μg of three recombinant SARS-CoV-2 antigens (nucleoprotein, spike subunit 1 and spike subunit 2) were used to capture specific serum antibodies, whereas a recombinant human protein (O^6^-methylguanine DNA methyltransferase) was used as a control antigen in the assay. Distinct MagPlex microsphere sets (Luminex Corp) were respectively coupled to viral or control antigens using the amine coupling kit (Bio-Rad Laboratories) according to manufacturers’ instructions. This three microsphere immunoassays (MIA) were developed and provided by Institut Pasteur, Paris. The MIA procedure was performed by incubating the serum samples (50 μl), diluted 1:400 in assay buffer (PBS-1% BSA-0.05% Tween 20), with the mixture of antigen-coated bead sets (about 1250 beads of each type) protected from the light on an orbital shaker at 700 rpm for 30 min. After washing, 50 μl of biotinylated protein A and biotinylated protein G (Thermo Fisher Scientific) at a 4 μg/ml each in assay buffer were transferred to each well and incubated on an orbital shaker for 30 min at 700 rpm in the dark. After washing, the beads were incubated for 10min at 700 rpm in the dark with 50 μl of Streptavidin-R-Phycoerythrin (Life technologies) diluted to 4 μg/mL in assay buffer. After washing, beads were resuspended in 100 μl of assay buffer. Measurements were performed using a Magpix instrument (Luminex), at least 100 events were read for each bead set and binding events were displayed as median fluorescence intensities (MFI). Relative Fluorescence Intensities (RFI) were calculated for each sample by dividing the MFI signal measured for the antigen-coated microsphere sets by the MFI signal obtained for the control microsphere set, to account for nonspecific binding of antibodies to beads. Specific seropositivity cut-off values for each antigen were set at three standard deviations above the mean RFI of the 37 dog (from France and Gabon) and 14 cat samples (from France) from the control group sampled before 2019. Based on the prepandemic population, MIA specificity was set at 97,3% for dogs and 100% for cats.

### Statistical analyses

Fisher’s exact test was used to analyze differences in antibody detection between the COVID19+ household group and the unknown status household group, as well as tests comparing cats and dogs in COVID-19+ households.

## Acknowledgements

We are grateful to the pet owners for giving us their permission to sample their pets. We thank all veterinarians that helped us with sampling, particularly Corentin Buisson, Julie Roggy and Maud Walliang for their important contribution to the sample collection and database. We thank Dr Thierry Buronfosse for the kind gift of pre-pandemic sera. We also thank Kurt McKean for English editing of the manuscript (https://octopusediting.com/). The study was funded by the French National Agency for Research (ANR-14-EBOL-003-01; Ebola virus in central Africa: genetic diversity, modalities of cross-species transmission and risk assessment of outbreak emergence, EBOFAQ), by IDEXLYON project of Université de Lyon as part of the “Programme Investissements d’Avenir” (ANR-16-IDEX-0005). This work was supported by French state funds managed by the Agence Nationale de la Recherche (ANR) within Programme Investissements d’Avenir, Institut Hospitalo-Universitaire FOReSIGHT (ANR-18-IAHU-0001) and Institut de Recherche pour le Développement (IRD).

## Author contributions

V.L. and E.L. conceived the study.

M.F., B.R., E.K., P.B., E.R., O.V., S.D., B.B., J.V., A.K., V.L. and E.L. designed and performed the experiments.

M.F., B.R, E.K., A.K., V.L, E.L. designed the work

All authors analyzed the data and interpreted and discussed the results.

M.F., V.L. and E.L. wrote the manuscript with input from all authors.

## Competing Interests

The authors declare no competing interests.

## Notes

### Competing Interest Statement

The authors have declared no competing interest.

