## Supplementary tables 1-2 for "High prevalence of SARS-CoV-2 antibodies in pets from COVID-19+ households"

**Supplementary Table 1**. Summary of pet seropositivity according to the number of positive tests and population.

| | Combination of the 4 serologies | | | | | |
|---|---|---|---|---|---|---|
| | 1 test MIA | 2 tests MIA | 3 tests MIA | 1 test MIA + Seroneutralization | 2 tests MIA + Seroneutralization | 3 MIA + Seroneutralization |
| Dogs COVID-19+ household | 2 (15,4%) | 1 (7,7%) | 0 | 2 (15,4%) | 0 | 0 |
| Cats COVID-19+ household | 6 (17,6%) | 6 (17,6%) | 2 (5,9%) | 0 | 1 (2,9%) | 5 (14,7%) |
| Dogs unknown status household | 1 (4,5%) | 1(4,5%) | 0 | 0 | 0 | 0 |
| Cats unknown status household | 2 (12,5%) | 1 (6,3%) | 1 (6,3%) | 0 | 0 | 0 |

**Supplementary Table 2.** Summary of pet seropositivity according to the serologic test and population

| | Serologic test | | | |
|---|---|---|---|---|
| | N | S1 | S2 | Seroneutralization |
| Dogs COVID-19+ household | 2 | 3 | 1 | 2 |
| Cats COVID-19+ household | 11 | 14 | 16 | 6 |
| Dogs unknown status household | 0 | 2 | 1 | 0 |
| Cats unknown status household | 2 | 2 | 3 | 0 |
